# Multiple Shoot Regeneration from Excised Embryonic Axis Explants of Jute, *Corchorus capsularis* cv. JRC321

**DOI:** 10.1101/2021.01.09.426027

**Authors:** Alpana Joshi, Subrata K. Das, Dipak Das

## Abstract

Interspecific hybrids development by traditional breeding is challenging because of the incompatibility between two cultivated species of genus *Corchorus*. Under this condition, plant genetic transformation is an alternative tool for the improvement of jute cultivar. In the present study, *in vitro* multiple shoot regeneration protocol has been developed in jute (*Corchorus capsularis* cv. JRC 321) from excised embryonic axis as an explant by removing cotyledons, radicle, and tip of epicotyl. The explant is cultured in medium SMM1 supplemented with BAP (1.0 mg/L) for one week and sub-cultured in SMM6 containing IBA (0.5 mg/L) and BAP (0.2 mg/L). Direct adventitious shoot bud formation was observed from cotyledonary nodes and the adjacent region of excised embryonic axis. Interestingly, shoot bud formation was reduced when excised explants with intact epicotyl were cultured in the media combination (SMM1+SMM6) indicating the presence of apical dominance might be the cause of fewer shoot bud formation. While, hypocotyl explants did not produce any shoot in this media combination because nodal meristem was removed. However, the shoot buds developed from the excised embryonic axis were differentiated successfully into multiple shoots and showed further elongation in SMM6 which ultimately rooted in MS-B5 medium (+0.5 mg/L IBA). The plants were transferred successfully in the glasshouse conditions and appeared phenotypically normal.

## Introduction

Jute is an important fiber crop having commercial significance in the production of diversified value-added products. It belongs to the genus *Corchorus* which comprises over 100 species and only two are cultivated, *C. capsularis* and *C. olitorius*^1^. Jute fiber is natural and biodegradable that can be used as an alternative to plastic or other synthetic materials. It is ligno-cellulosic in nature and the presence of lignin in the fiber adversely affects its quality. Interspecific hybridization is a method can be used to combine useful characters of the two species in a single genotype like improved fiber quality with altered or reduced lignin content, better yield, higher retting of fiber, and resistance against biotic and abiotic stress. However, the main problem associated with Jute variety is a strong sexual incompatibility between the two cultivated species hindered natural or conventional breeding to create interspecific hybrids^2^.

The other possible ways to improve the fiber crop are protoplast fusion of the two species having unique characters and genetic transformation^3–8^. Limited success in obtaining embryo like structures from protoplasts was reported^9^. Genetic transformation can be done by utilizing the organogenesis as well as somatic embryogenesis potential of preformed regenerable structures such as cotyledonary nodes, cotyledonary petioles and epicotyl tip. Stable genetic transformation of *C. capsularis* by particle bombardment using shoot tip explants has been reported^10^. *In-vitro* regeneration using cotyledon and petiole attached cotyledon for different cultivars from both the species of *Corchorus* was reported^11–16^. Limited regeneration from leaf explants of *C. olitorius* was observed^4^. Somatic embryogenesis from cotyledon and hypocotyl protoplast derived calluses in *C. capsularis* was reported^7^. However, a very limited or no shoot regeneration was documented.

But, due to the highly recalcitrant nature and genotype dependent of the two cultivated species of *Corchorus* in tissue culture, major improvement of this crop by means of genetic transformation could not be achieved yet^17^. However, standardization of preliminary transformation and regeneration using cotyledonary petiole explants of jute (JRC321) was documented^18^. Also, limited plantlets (2-3 shoots) regeneration using shoot meristem-tip and epicotyl explants of seedling, and immature shoot bud explant after *Agrobacterium* transformation^19^. High frequency of shoot regeneration is required for efficient transformation. Therefore, development of direct multiple shooting from induced meristematic zone using tissue culture method need to be more focused to encounter the transformation problem.

In the present study, we successfully achieved multiple shoot regeneration from excised embryonic axis from germinating seeds of *Corchorus capsularis* cv. JRC 321.

## Materials and Methods

### Ex-plant preparation for multiple shooting

Mature seeds of *C. capsularis* cv. JRC 321 were used as the experimental material for regeneration. The seeds were surface sterilized with 0.2% mercuric chloride solution for 5 mins and subsequently washed (five times) with sterile water in a laminar flow bench. The seeds were soaked in sterile water for overnight at dark condition and placed at 30°C on growth-regulator free half strength of MS^20^ basal medium with MS salts and B5 vitamins^21^, 1% (w/v) sucrose and 0.8% agar (HiMedia, Mumbai, India) for germination. Seed coats were removed by scalpel, cotyledonary leaves and root axis of 2, 3, 4, and, 5 days of germinating seeds were removed and cut transversely to make little incision on the tip of epicotyl as indicated with dotted lines (Fig. 1A). The excised embryonic axis of different stages was used as explants. The explants were placed on shoot multiplication medium (SMM) for shoot regeneration.

**Figure 1.**
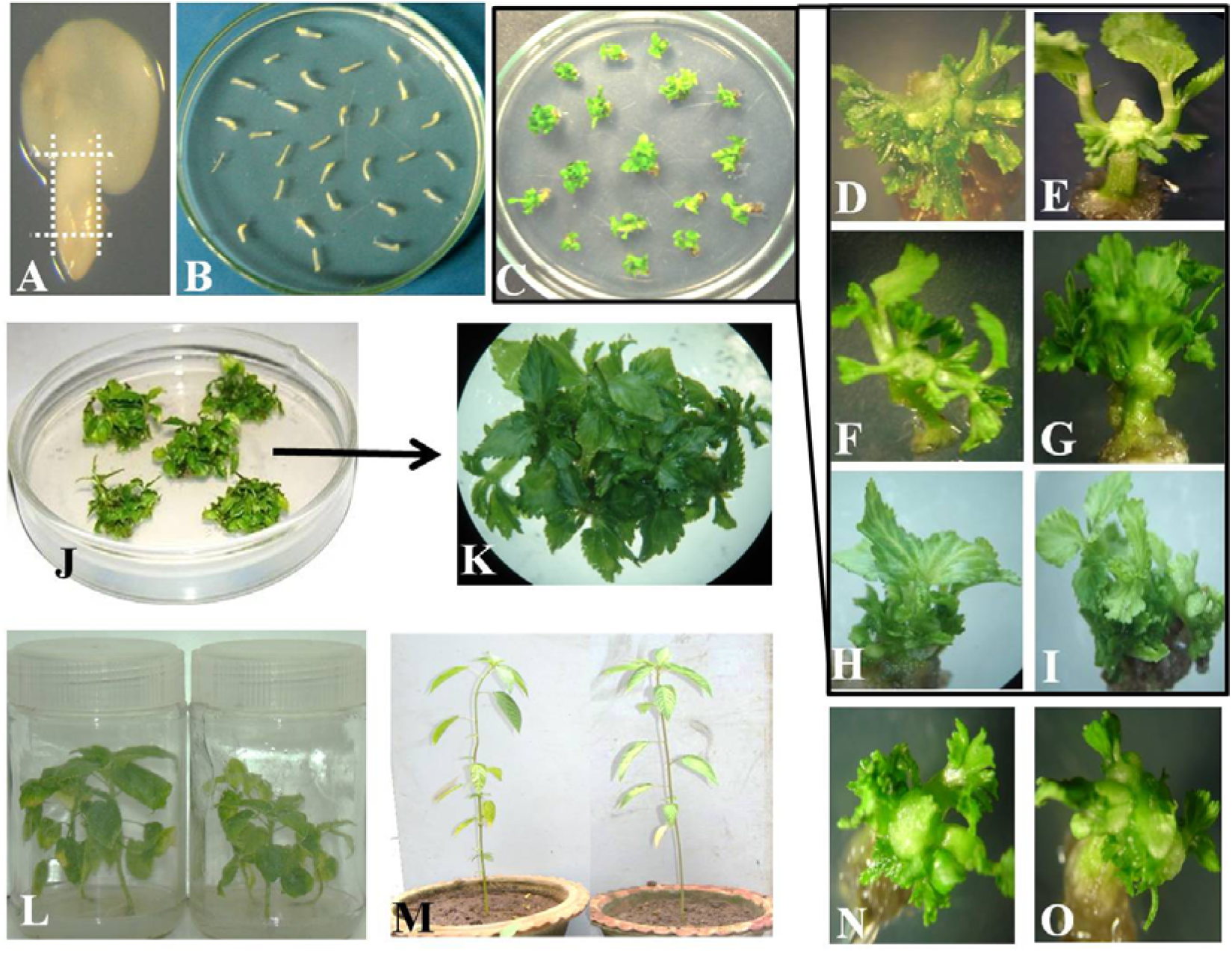
Regeneration of Multiple Shoots from Excised Embryonic Axis of Jute (*Corchorus capsularis* cv JRC 321): **(A)** Embryo from 2 days old germinated seed and the dotted lines indicate 4 cuts were made to prepare explant; **(B)** Explants on the SMM1 medium containing 1.0mg/L BAP for 1 week; **(C)** Adventitious shoot bud regeneration from excised embryonic explants after 7-10 days of transfer on SMM6 medium (0.5mg/L IBA and 0.2 mg/L BAP); (**D–I)** 5X magnified picture (stereo-zoom microscopy) of regenerated explants with adventitious shoot buds shown in Figure 1C; **(J)** Multiple shoot regeneration in petri plate after three subcultures in SMM6 medium; **(K)** 5X magnified picture (stereo-zoom microscopy) of regenerated shoots from Figure 1J; **(L)** Plantlets were transferred from petri plates to culture bottle containing MS basal medium supplemented with 0.5mg/L IBA for proper rooting and elongation; **(M)** Regenerated mature plantlets; **(N & O)** Morphology of the malformed multiple shoots under stereo-zoom microscope showing stunted growth at higher concentration of BAP. No differentiations were observed among the developmental stages of the initiated shoot buds.

To know the effect of incisions at different points of embryonic axis for multiple shoot regeneration, three types of explants were prepared. Type-I explant with 4 cuts on embryo removing radical, cotyledons and tip of epicotyls, Type-II with 3 cuts on embryo removing radical and cotyledons keeping epicotyl tip intact, and Type-III with 4 cuts on embryo removing radical, cotyledons and epicotyls keeping only hypocotyls of embryo axis (Fig. 2 A, C & G).

**Figure 2.**
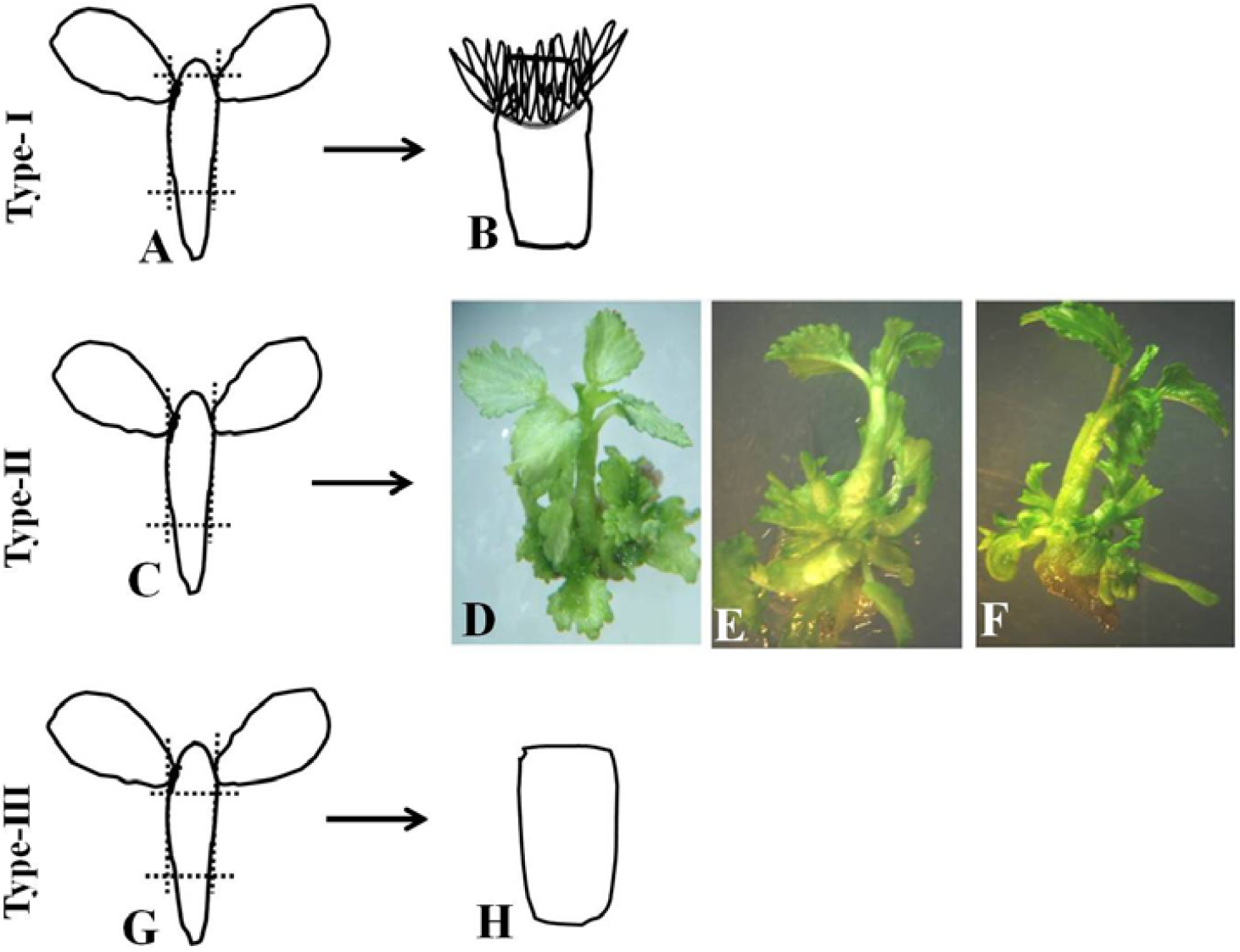
Diagrammatic Representation of Explants Preparation and Effect of Explants on Shoot Bud Regeneration Efficiency: **(A)** Showing preparation of Type-I explants: dotted lines indicate 4 cuts on embryo removing radical, cotyledons and tip of epicotyl; **(B)** Diagram showing adventitious shoot duds formation from excised embryo axis; **(C)** Type-II explants prepared by (dotted lines) 3 cuts on embryo, removing radical and cotyledons; **(D-F)** Pictures showing shoot buds with single apical shoot indicates the formation of 7-10 shoot buds without suppressing apical dominance; **(G)** Type-III explants (Hypocotyl), dotted lines indicate 4 cuts on embryo by removing radical, cotyledons and epicotyls; **(H)** Diagram showing no adventitious shoot buds regeneration from Type-III explants.

### Media preparation and growth condition

Multiple shoot regeneration in jute explants was investigated in the presence of different concentration of auxins (2, 4-D, IAA, IBA and NAA) and cytokinins (kinetin, BAP and TDZ) in MS basal medium (MS salts + B5 vitamins + 2% sucrose + 500 mg/L glutamine). The explants were placed on MS basal medium supplemented with different combination of growth regulators and incubated at 27°C for 16/8 h light/dark photoperiod for the duration of 6 weeks. Observations were recorded periodically. The explants were sub-cultured to fresh media in every week of culture. The most responsive medium was selected in terms of good multiple shoot regeneration. When the explants started to multiply, well grown shoots were separated with the help of a sterile scalpel under the hood and put in the same media for further multiplication. All media were sterilized at 120°C for 15 mins and pH of the medium was adjusted to 5.8.

### Rooting and hardening of plantlet

The multiplied explants were carefully separated with the help of a sterile scalpel and then placed in the MS basal medium without growth regulators for two weeks. After that, the plantlets were transferred into the MS basal medium supplemented with 0.5mg/L IBA for proper rooting and elongation in the culture bottle^13^.

Plantlets were removed from culture bottle and washed to remove the medium with it and placed to sterile vermiculite and incubated under culture room conditions. The acclimatized plants (80%) from the culture room conditions were carefully transferred to pots filled with soil and manure. All regenerated plants appeared normal with respect to morphology, growth and fruit set.

## Results & Discussion

### Effect of age of explants on efficiency of multiple shoot induction from excised embryonic axis

Excised embryonic axes collected from different stages of germinated seeds of jute were used as explants to investigate the effect of age on *in-vitro* shoot regeneration. The experiment was performed initially in SMM6 (0.5 mg/L IBA and 0.2 mg/L BAP) based on previous report^10^. Maximum number of multiple shooting was observed from the embryonic axis of 2 days old germinating seeds with an average regeneration frequency of 10 shoots per explant (Table 1) whereas low regeneration frequency was observed in older ages of germinated seeds (4-5 days old seeds). It showed that cotyledons from 2 days old germinating seeds exhibited a high level of plant regeneration ability when cultured in liquid BM medium in the presence of NAA and BAP^13^. Further experiment was done on explant from 2 days old germinated seeds.

**Table 1:** Effect of Explants Age on the Efficiency of Multiple Shoot Induction in SMM6.

| **Age of explants**<br>**(Shoot primordia)** | **No. of explants**<br>**inoculated** | **No. of explants showing**<br>**multiplication** | **Average frequency of**<br>**shoots/explant** |
| --- | --- | --- | --- |
| 2 days old | 25 | 20 | 10 |
| 3 days old | 25 | 15 | 6 |
| 4 days old | 25 | 12 | 3 |
| 5 days old | 25 | 12 | 3 |

### Effect of growth regulators on efficiency of multiple shoot induction

Excised embryonic axes were cultured on MS basal medium supplemented with different combination and concentration of auxins and cytokinins, designated as SMM (shooting multiplication medium) media to study the effect of growth regulators on multiple shoot regeneration of 2 days old germinating seeds. The basal medium without plant growth regulator was taken as a control. The combination of growth regulators was used *viz*., 2,4-D+kinetin, NAA+BAP, IAA+BAP, IAA+TDZ, NAA+TDZ and, IBA+BAP. All tested combinations except IBA+BAP were failed to induce the generation of multiple shoots from the explants. However, the combination of these hormones (2,4-D+kinetin, NAA+BAP, IAA+BAP, IAA+TDZ, NAA+TDZ) produced large, hard, green callus at the base of explants and regenerate few shoots (0-3 shoots/explant). Multiple shooting with varying frequencies was observed in presence of different combinations of IBA+BAP (SMM1-SMM6, Table 2). The higher frequency was observed on the basal media (SMM6) containing 0.5 mg/L of IBA and 0.2 mg/L BAP compared to other media combinations. Higher concentration and prolonged exposure with BAP enhanced adventitious shoot bud initiation from the explants but the regenerated shoots were found to be stunted in growth (Fig. 1 N & O). On the other hand, lowering the concentration of BAP produced less number of shoots but no malformation was observed. The best result was observed when embryonic axes were cultured in SMM1 (1.0mg/L BAP) for one week and then transferred these explants in SMM6 (0.5mg/L IBA + 0.2mg/L BAP) that showed highest frequency of shooting within 40-50 days. However, the initiation of direct multiple shooting was observed within 7-10 days in the medium SMM6 (Fig. 1C-I). The branching of multiplied shoots was being continued when the explants were cultured in the SMM6 at least for two weeks. Thus, shoots generated in combination of SMM1 and SMM6 medium was found to be effective for generation of plantlets.

**Table 2:** Effect of BAP and IBA on efficiency of multiple shoot induction.

| **Explant inoculated in different media** | **No. of the explants inoculated** | **Percentages of explants showing shoot multiplication** | **Average frequency of normal shoots/explant** |
| --- | --- | --- | --- |
| SMM1 (MS+1mg/L BAP) | 100 | 33 | 0 |
| SMM2 (MS+0.5mg/L BAP) | 100 | 50 | 1 |
| SMM3 (MS+0.2mg/L BAP) | 100 | 78 | 3 |
| SMM4 (MS+1mg/L BAP+1mg/L IBA) | 100 | 40 | 0 |
| SMM5 (MS+0.5mg/L BAP+ 0.5mg/L IBA) | 100 | 57 | 3 |
| SMM6(MS+0.2mg/L BAP+0.5mg/L IBA) | 100 | 93 | 10 |
| SMM1(1<sup>st</sup> week)+ SMM6(2<sup>nd</sup> -8<sup>th</sup> week) | 100 | 85 | 24 |

**Table 3:**
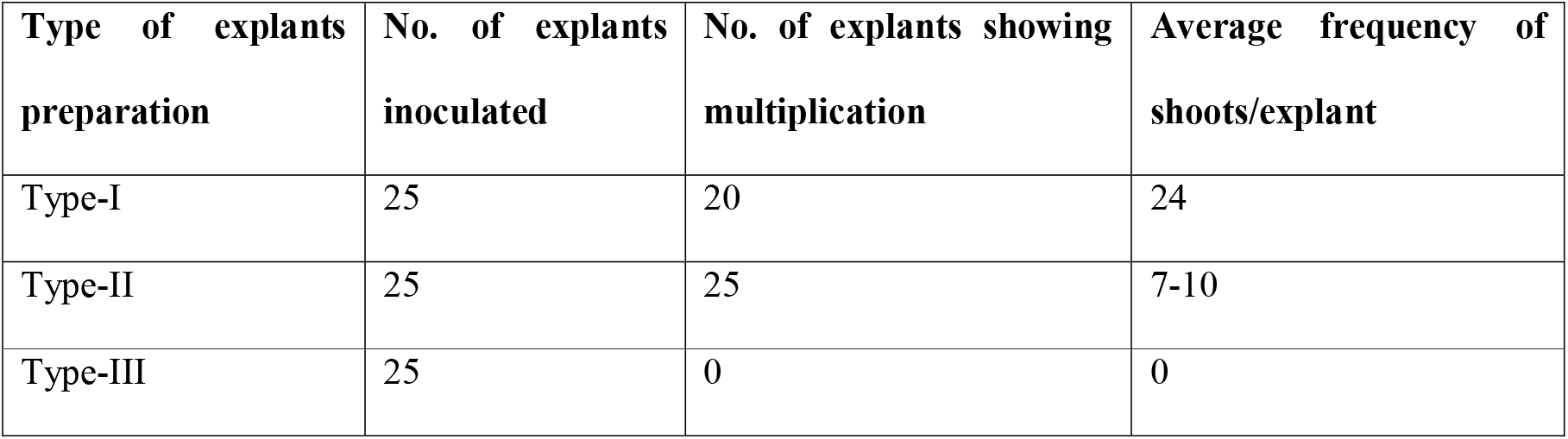
Effect of explants preparation on multiple shoot regeneration in SMM1 + SMM6 media combination.

BAP is the most effective kinetin in promoting multiple shoot regeneration in various plants including jute. Best multiple shooting from petiole attached cotyledon explants was found when they were cultured on MS containing 0.2 mg/L BAP and 1.0 mg/L IAA in CVL-1 and D-154 of *C. capsularis*^16^. On the other hand, CVE-3 showed best response on MS with 2.5 mg/L BAP and 0.5 mg/L NAA. BAP induce multiple shoot production by stimulating rapid cell division^22,23^. Shoot production was improved in response to BAP application^24^. In the present study, the exposure of explants with BAP (1.0mg/L) in SMM1 medium for one week might enhance the meristematic activity adjacent to cotyledonary nodal region and induce initiation of adventitious shoot bud formation. Subsequent treatment of explants by reducing the BAP concentration to 0.2mg/L and inclusion of IBA (0.5mg/L) in the medium (SMM6) enhance differentiation of shoot buds towards multiple shoots and further elongation of those shoots was observed. However, the higher concentration and prolonged exposure of BAP caused inhibitory effect on the elongation of adventitious shoot buds. Our result was supported by the earlier finding where BAP effectively induced shoot bud formation at higher concentrations but inhibited further development and growth of shoot buds in *Vigna radiata*^25^. The author also stated that lowering BAP concentration at late stage and inclusion of auxin efficiently induced multiple shooting from cotyledonary node culture in *Vigna radiata*. Other studies revealed that direct shoot buds are originated from induced meristematic region of seedling in *Withania somnifera* and *Glycine max*, respectively^26,27^.

### Effect of incision in shoot regeneration from different explants

The present finding suggest that the explant preparation is very important for shoot regeneration. Shoot regeneration experiment with three type of embryonic axis was conducted to study the effect of incision at different point of explants isolated from 2 days old germinated seeds. Explants (Type-I, II and, III) was cultured with SMM1 for one week and subsequently sub-cultured in SMM6 media, only Type-I explants showed adventitious shoot buds whereas few shoot buds with main apical shoot in Type-II explants and, no buds or shoot in Type-III explants was observed (Fig. 2). This study indicates that Type-I explant able to regenerate into multiple shoots as these explants has incision on the tip of epicotyl which suppressed the apical dominance. Moreover, the exposure of Type-I explants with SMM1 medium (1mg/L BAP) possibly stimulates the formation of adventitious shoot buds from cotyledonary node and its adjacent region of embryonic axis.

In contrast, the shoot bud formation was reduced dramatically in Type-II explants in the (SMM1 + SMM6) media combination indicating the presence of apical dominance might decrease shoot bud formation. Whereas the higher frequency of shoot regeneration in Type-I explants might be due to increased meristematic activity of cotyledonary nodal and adjacent region of embryonic axis. Although, shoot bud was not induced in Type-III explants (contain only hypocotyl region), as nodal meristem was removed. Therefore, the current protocol suggests that Type-I incision and sequential SMM media treatment resulted induction and proliferation of multiple shoots from excised embryonic axis. Rozali et al also used the incision technique for explant preparation and showed that micro shoots were highly induced when the young shoot bud explants, incised prior subculture^28^. Moreover, the wounding near to meristematic zone not only induces adventitious shoot bud formation but also essential for *Agrobacterium* mediated transformation. This multiple shooting protocol may be effectively used for the improvement of Jute cultivar through *Agrobacterium*-mediated or particle bombardment methods of genetic transformation.

## Acknowledgement

The research work is carried out at School of Biological Engineering and Life Science, Shobhit Institute of Engineering & Technology (Deemed to be University, approved), NH-58, Modipuram, Meerut-250110 for providing facilities.

